# The molecules lost NEW COMPOUNDS BINDING AT THE HISTAMINE SITE OF THE NMDA RECEPTOR (NMDA_(HA)_R)

**DOI:** 10.1101/2024.03.08.583891

**Authors:** V. Armand

## Abstract

NMDA receptor ligands have been the target of intensive research for the treatment of psychotic diseases and central nervous system diseases. Our group published the characterization of the NMDA receptor histamine site. We developed modulators for this site, about 500 drugs were tested and we finally have a partial agonist FUBn293 with a nanomolar affinity and also an antagonist ST-579 with a nanomolar affinity. We suggest that agonists at the (NMDA_(HA)_R) constitute an innovative class of antipsychotics for the treatment of schizophrenia and other neurological or psychiatric disorders.

---

In this article, I will relate more than 20 years of research on a series of molecules that would have been forgotten without this short article summarizing the activity structure of this one. This work was done by many researchers and I will list them here in the hope that I will not forget one, here they are:

V. Armand, J.-M. Arrang, C. Bayard, A. Burban, R. Faucard, C. R. Ganellin, S. Graßmann, P. P. Griffin, H. Kubas, I. Nuss, U. Reichert, W. Schunack, J.-C. Schwartz and H. Stark.

During this period, several science theses were written:

*<u>For the pharmacological results</u>*

in 1999 Cécile Bayard in Paris,

**Caractérisation du site de liaison de l’histamine du recepteur NMDA**.

in 2004 Raphaêl Faucard in Paris,

**Caractérisation pharmacologique et fonctionnelle du site histamine, modulateur du récepteur NMDA**.

in 2009 Aude Burban in Paris,

**Modulation du récepteur NMDA du glutamate par l’histamine: intérêt dans la schizophrénie**.

*<u>For the compounds formula and method for preparing</u>*

in 2000 Sven. Graßmann in Berlin,

**Synthese und Pharmakologie von Liganden der Histaminbindungsstelle des N-Methyl-D-aspartat-Rezeptors**

in 2000 Ulrich Reichert in Berlin,

**Heterocyclische Alkanamine als Modulatoren der Histamin-Bindungsstelle des NMDA-Rezeptors: Synthese, Analytik und Struktur-Wirkungsbeziehungen**.

in 2004 Isabelle Nuss in Berlin,

**Heterobicyclische Alkanamine als Liganden einer modulatorischen Bindungsstelle des N-Methyl-D-aspartat-Rezeptors: Synthese, Analytik und Struktur-Wirkungsbeziehungen**.

in 2005 Perry Paige. Griffin in Frankfurt am Main,

**Neue Liganden einer modulatorischen Bindungsstelle an NMDA-Rezeptoren: Synthese, Analytik und Struktur-Wirkungsbeziehungen**.

in 2007 Holger Kubas in Frankfurt am Main,

**Substituierte Benzothiazole als allosterische Modulatoren an NMDA-Rezeptoren**.

There have also been communications on the subject;

U. Reichert, S. Graßmann, C. Bayard, J.-M. Arrang, J.-C. Schwartz, H. Stark und W. Schunack.

**Heterocyclische Alkanamine als positive Modulatoren des NMDA-Rezeptors.Deutsche Pharmazeutische Gesellschaft: Landesgruppe Berlin-Brandenburg: Der wissenschaftliche Nachwuchs stellt sich vor. Berlin (5. Juli 1999), Abstractbuch**.

S. Graßmann, U. Reichert, C. Bayard, J.-M. Arrang, J.-C. Schwartz, H. Stark und W. Schunack.

**Neue Leitstrukturen für Agonisten einer modulatorischen Bindungsstelle des NMDA-Rezeptors**.

**Deutsche Pharmazeutische Gesellschaft: Landesgruppe Berlin-Brandenburg: Der wissenschaftliche Nachwuchs stellt sich vor. Berlin (5. Juli 1999), Abstractbuch.**

I Nuss, U. Reichert, S. Graßmann, R. Faucard, C. Bayard, J.-M. Arrang, J.-C. Schwartz, H.Stark, and W. Schunack.

**Synthesis and Pharmacology of Novel Potent Ligands at the Histamine Binding Site of the NMDA Receptor. 3rd European Graduate Student Meeting of the German Pharmaceutical Society (DphG), Frankfurt am Main (23.-25. Februar 2001). Arch. Pharm. Pharm. Med. Chem. 2001, 334 (Suppl. 1), 23 (A-65)**.

I. Nuss, S. Graßmann, R. Faucard, J.-M. Arrang, J.-C. Schwartz, H. Stark und W. Schunack.

**Synthese und Pharmakologie von neuen potenten Liganden der Histamin-Bindungsstelle des NMDA-Rezeptors. Deutsche Pharmazeutische Gesellschaft: Landesgruppe Berlin-Brandenburg: Der wissenschaftliche Nachwuchs stellt sich vor. Berlin (12. Juli 2001), Abstractbuch**.

S. Graßmann, C. Bayard, J.-M. Arrang, C. R. Ganellin, J.-C. Schwartz, H. Stark, and W. Schunack.

**In Vitro Investigation on Novel Potent Antagonists of a Modulatory Binding Site of NMDA Receptors. Jahrestagung der Deutschen Pharmazeutischen Gesellschaft (DPhG) „Die wissenschaftliche Pharmazie im neuen Jahrtausend – Trends, Entwicklungen, Highlights“, Halle (Saale) (10.-13. Oktober 2001). Arch. Pharm. Pharm. Med. Chem. 2001, 334 (Suppl. 2), 18 (V1-17).**

P. P. Griffin, S. Graßmann, C. Bayard, J.-M. Arrang, J.-C. Schwartz, W. Schunack, and H.Stark.

**Histamine Derivatives as Potential NMDA Receptor Ligands. Deutsche Pharmazeutische Gesellschaft (DPhG) – Jahrestagung 2002, Berlin (9.-12. Oktober 2002), Poster. Arch. Pharm. Pharm. Med. Chem. 2002, 335 (Suppl. 1), 81 (P:C36)**.

I. Nuss, S. Graßmann, R. Faucard, J.-C. Schwartz, J.-M. Arrang, H. Stark, and W. Schunack.

**Bicyclic Alkanamines as Highly Potent Ligands of a Modulatory Binding Site of NMDA Receptors. Deutsche Pharmazeutische Gesellschaft (DPhG) – Jahrestagung 2002, Berlin (9.-12.Oktober 2002), Koautor. Arch. Pharm. Pharm. Med. Chem. 2002, 335 (Suppl. 1), 48 (V:C16)**.

P. P. Griffin, S. Graßmann, C. Bayard, J.-M. Arrang, J.-C. Schwartz, W. Schunack, and H. Stark

**Cytoprotective Histamine Derivatives as NMDA Receptor Modulators. ZAFES Kick-Off Symposium “Lipid Signaling”, Frankfurt am Main (14. Oktober 2004)**.

The project was stopped more than 10 years ago because the patent on these molecules entitled; **NMDA RECEPTOR HISTAMINE SITE MODULATORS FOR USE IN THE TREATMENT OF CENTRAL NERVOUS SYSTEM DISEASES** was not accepted, some of the data had been released to the public, confer to the symposium.

So these are the molecules I think it’s a shame to forget them for good.

## Introduction

The effects of histamine are mediated by four G protein-coupled receptors (H_1_, H_2_, H_3_ and H_4_). In the brain, histamine also binds to the histamine site (NMDA_(HA)_R) of the N-methyl-D-aspartate receptor (NMDAR) (1). Histamine potentiates NMDA currents in isolated (2), and cultured (3) hippocampal neurons and this effect requires NMDARs containing NR1 variants lacking exon 5 with NR2B subunits (4, 5). This potentiation is inversely related to the concentration of glycine (2, 6) and is reproduced by tele-methylhistamine (tele-MeHA), the catabolite of histamine in the brain (3, 4–6). Histamine also binds to NMDA_(HA)_R to potentiate NMDA-induced [^3^H]noradrenaline release from hippocampal synaptosomes(6). Histamine potentiates N-methyl-D-aspartate receptors by interacting with an allosteric site distinct from the polyamine binding site (6)

## Materials and Methods

[^3^H] noradrenaline release from hippocampal synaptosomes.

A crude synaptosomal fraction was prepared as described previously with minor modifications (7).Adult male Wistar rats (200-250g) were killed by decapitation. The hippocampus wa rapidly removed and homogenized (Potter Elvehjem glass; eight up-down strokes) in 40 volumes of 0,32 M sucrose. The homogenate was first cecntrifuged (100g for 10 min) to remove nuclei and cellular debris. Synaptosomes were isolated from the supernatrant by a second centrifugation (12,000 g for 20 min). The synaptosomal pellet was then suspended in modified Krebs-Ringer bicarbonate medium of the following composition: NaCl 120 mM; KCl 0.8 mM; KH2PO4,1.2 mM; CaCl2 1.3 mM; MgSO4 1,2 mM; NaHCO3 27.5 mM; glucose, 10 mM; ascorbic acid 0.06 mM; EDTA 0.03 mM; gassed with 95% O2 and 5% CO2); pH 7.4.

Synaptosomes are then suspended in this same medium in the require volume of assay buffer abd are then incubated for 1 hour at 37°C in a rotary water bath, in an atmosphere of 95% 02 and 5% CO2, with [3H]noradrenaline (final concentration 30 nM, GE Healthcare, Buckingamshire, UK). During this incubation, synaptosomes are loaded with the labelled neurotransmitter. Then, the labelling of synaptosomes is followed by 4 washes with a Mg2+-free medium prewarmed at 37°C.

Synaptosomes are distributed in identical aliquots (200 µg of protein) in a final volume of 500 µL and incubated with NMDA (200µM), glycine (1 µM) and drugs to test in the presence of thioperamide, an H3 receptor antagonist in a saturating concentration (1 µM) to prevent the action of the heteroreceptor H3 modulating [^3^H] noradrenaline release in the system.(8) After 3 minutes of incubation at 37°C, reaction is stopped by immersion of tubes in ice-cold water, immediately followed by a centrifugation (14 000 × g, 10 sec). The amount of radioactivity released into each supernatant is finally determined by liquid scintillation using a β counter.

## Results

We have investigated new drugs, ligands of the histamine site of the NMDA receptor by using the model of the NMDA-mediated [^3^H]noradrenaline release from hippocampal synaptosomes.

Over the years, we have tested more than 500 molecules and identified agonists and antagonists of the histamine binding site on the NMDA receptor, we present the most remarkable molecules with structure activity relationships in two series of tables in supplementary datas with the agonist (table 1-9) and the antagonist (table 11-14)

On this response we identify some 2-aminobenzothiazole derivatives as potent agonists of the NMDA_(HA)_R. The FUB_n_7, is the lead compound, it was a full agonist with a micromolar potency. Optimisation of this lead was obtained with substitution of the aliphatic amino group, that led to agonists with a nanomolar agonist potency such as FUB_n_293(9,10).

The FUB_n_7,as lead coupound was first tested in various brain regions and the hippocampus give the best answer on [^3^H]noradrenaline release (Fig 1B), then we used several NMDAR antagonist on the FUB_n_7 and we observed a non competive inhibition (Fig 1C,D,E).

**Fig 1.**
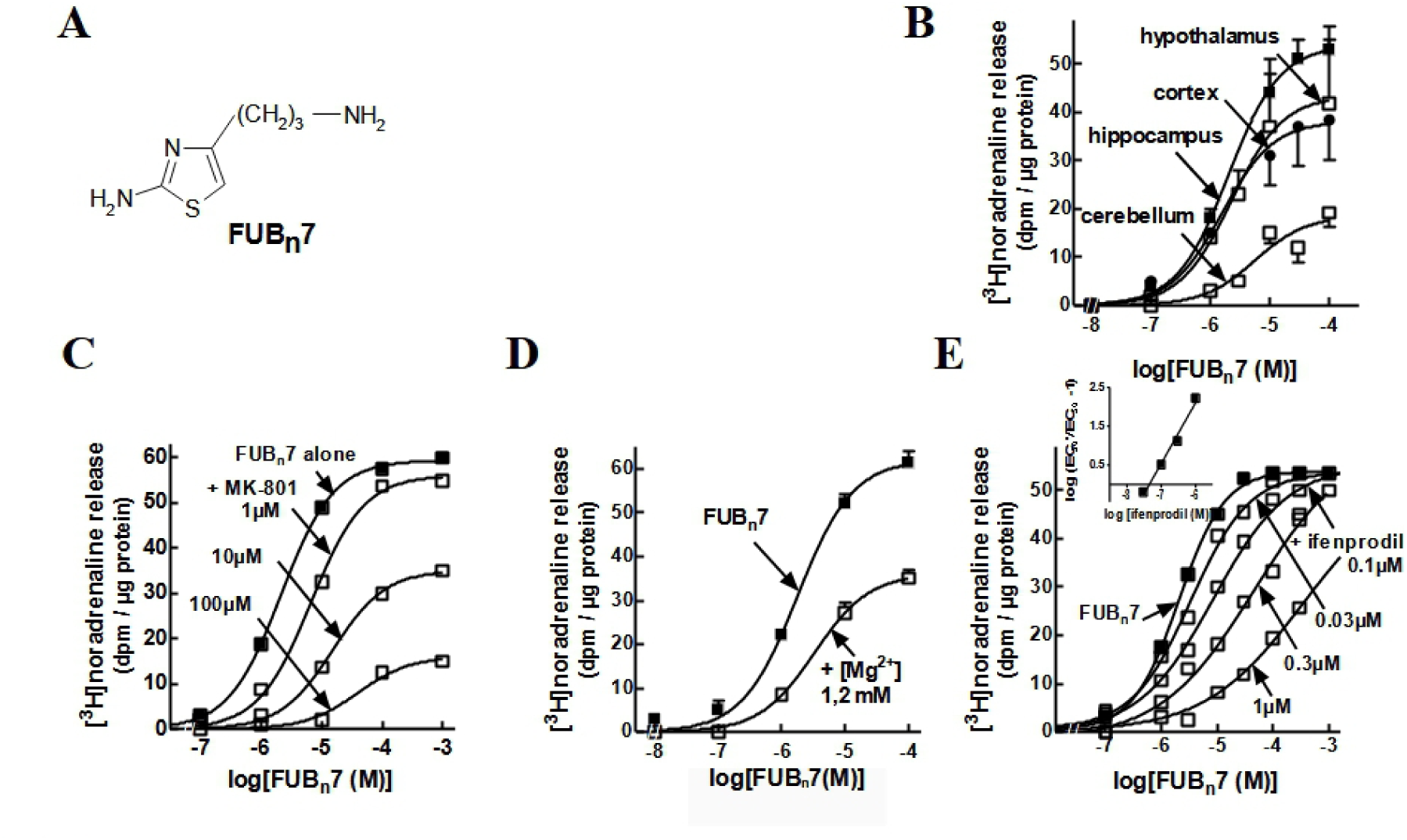
Effects of FUBn7 NMDA-induced [3H]noradrenaline release. **A**, Chemical structure of FUBn7.. **B**, Effect of FUBn7 on NMDA-induced [3H)noradrenaline release from synaptosomes of various rat brain regions. **C, D, E**, Effect of MK-801 (1-100 µM) (**C**), Mg2+ (1.2 mM) (**D**) and ifenprodil (0.03-1 µM) (**E**) on NMDA-induced [3H)noradrenaline release from hippocampal synaptosomes. Results are expressed as dprn/µg protein over [3H]noradrenaline release induced by NMDA (200µM) and glycine (1µM). Eachpoint represents the mean ± SEM of values obtained in 3-8 separate experiments.

FUB_n_7, the first lead compound obtained in this series, behaved as a full agonist with a micromolar potency (EC_50_ = 2.1 ± 0.1 µM). Its effect was antagonized by the NMDAR blockers MK-801 (Figure 1C) and by ifenprodil, the NR2B antagonist (Figure 1E).

As we previously reported (9,10), FUB_n_293 also potentiated NMDA-induced [^3^H]noradrenaline release from hippocampal synaptosomes (EC_50_ = 2,8 ± 1.8 nM). FUB_n_293 displayed a nanomolar agonist potency on NMDA-induced [^3^H]noradrenaline release from hippocampal synaptosomes. It was around 1000 fold more potent than FUB_n_7 and 25,000 fold more potent than histamine, but its maximal effect was 49 ± 6% that of FUB_n_7, suggesting that it behaved as a partial agonist on this response (Fig 2B).

**Fig 2.**
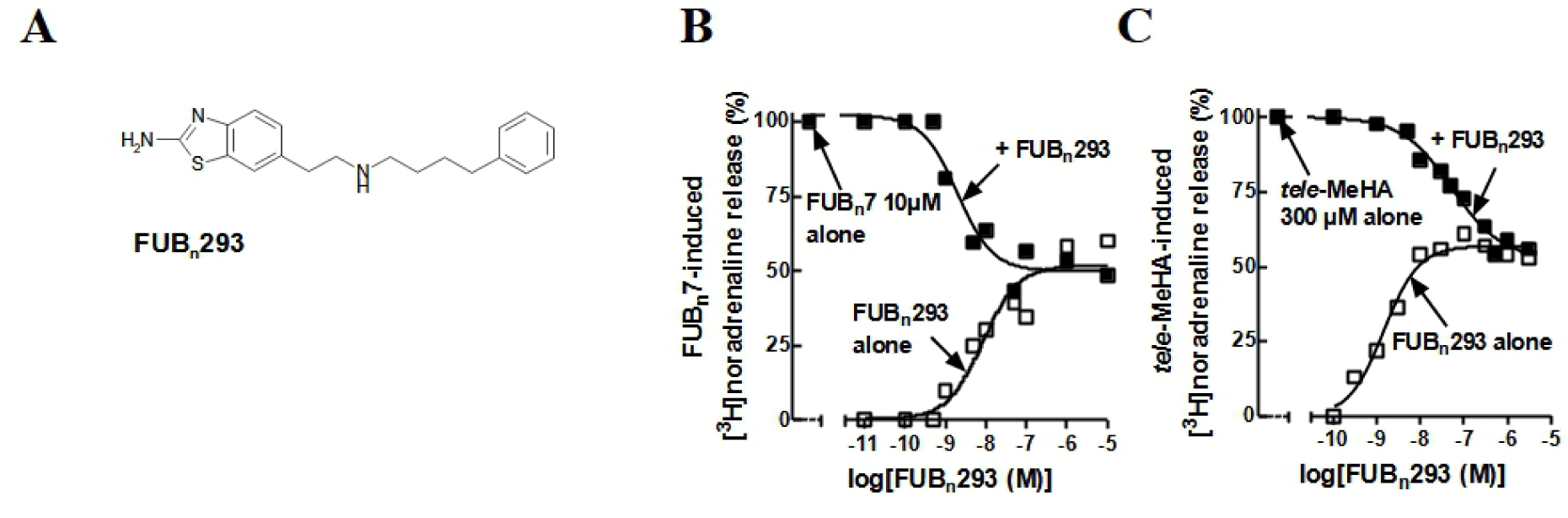
Effects of FUBn293 NMDA-induced [JH]noradrenaline release. **A**, Chemical structure of FUBn293. **B, C,** Effect of FUBn293 tested alone, or against FUBn7 **(B),** or tele-MeHA **(C),** on NMDA-induced [3H]noradrenaline release from hippocampal synaptosomes Results are expressed as per cent of the effect of FUBn7 **(B),** or tele-MeHA **(C)** (means of 6-16 determinations from 3-8 separate experiments).

In agreement, FUB_n_293 decreased in a concentration-dependent manner the sub-maximal effect of FUB_n_7. The maximal antagonism reached at the highest concentrations tested led to the same plateau as its maximal agonist effect (–50.2 ± 3.1% vs +50.9 ± 4.1% of FUB_n_7-induced release), and its *K*_i_ assuming a competitive antagonism of FUB_n_7 was in the same nanomolar range as its agonist potency (3.7 ± 1.4 nM) (Figure. 2B). A similar pattern and *K*_i_ value (7.6 ± 1.9 nM) was obtained when FUB_n_293 was opposed to *tele*-MeHA (Figure. 2C), which confirmed that FUB_n_7 and FUB_n_293 bind at the histamine site i.e. the NMDA_(HA)_R.

We obtain also antagonists, the best was the ST-579 (10), ST-579 was able to inhibite the potentiation of [^3^H]noradrenaline release induced by FUB_n_7 (Fig 3B) and FUB_n_293 (Fig 3C)

**Fig 3.**
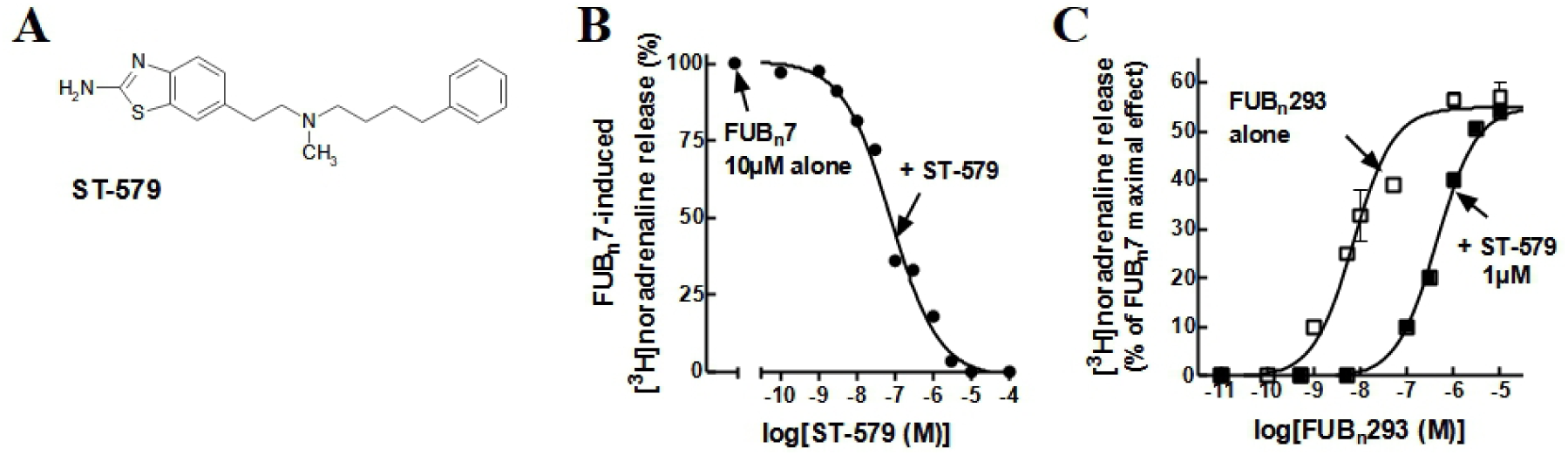
Effects of ST-579 on NMDA currents and NMDA-induced [JH]noradrenaline release. **A**, Chemical structure of ST-579 **BC** Inhibition by ST-579 of the potentiation of NMDA-induced [3H]noradrenaline release induced by FUBn7 **(B)** or FUBn293 **(C)** Results are expressed as dprn/µg protein over [3H]noradrenaline release induced by NMDA (200µM) and glycine (1µM) Results are means of 6-9 determinations from 3 separate experiments.

## Activity Structure

The activity structure of a part of the compounds is presented in tables.

## THE AGONIST

The tables give the intrinsic activity of the drugs compared to the FUB 7 and the EC 50 of the drugs in µM.

The first series describes agonists, in which the molecules are derived from the general formula around the lead compound FUBn 7 (Table 1 to Table 3)

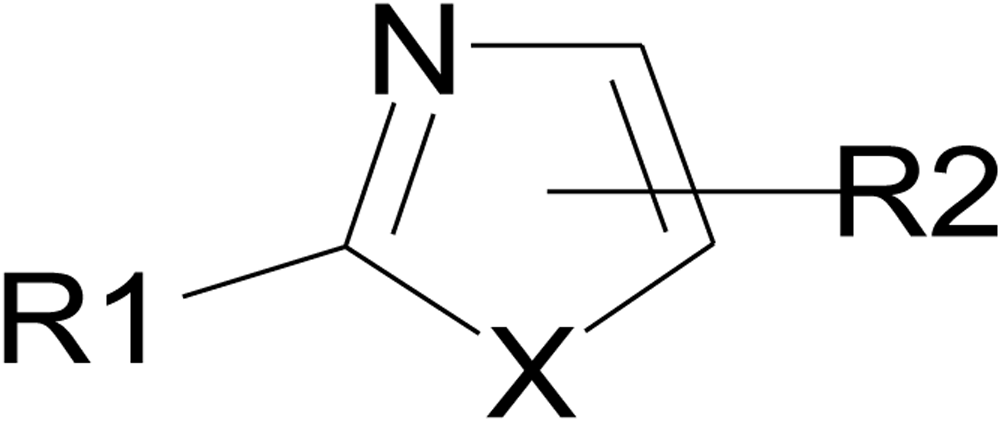

The second series describes agonists, in which the molecules are derivated from the general formula around the lead compound FUB 16 (derivated from R-1, 3-benzothiazol-2-amine) and lead to a first best compound FUB 247 and then to the FUBn293 the most potent drug realized (Table 4 to Table 9).

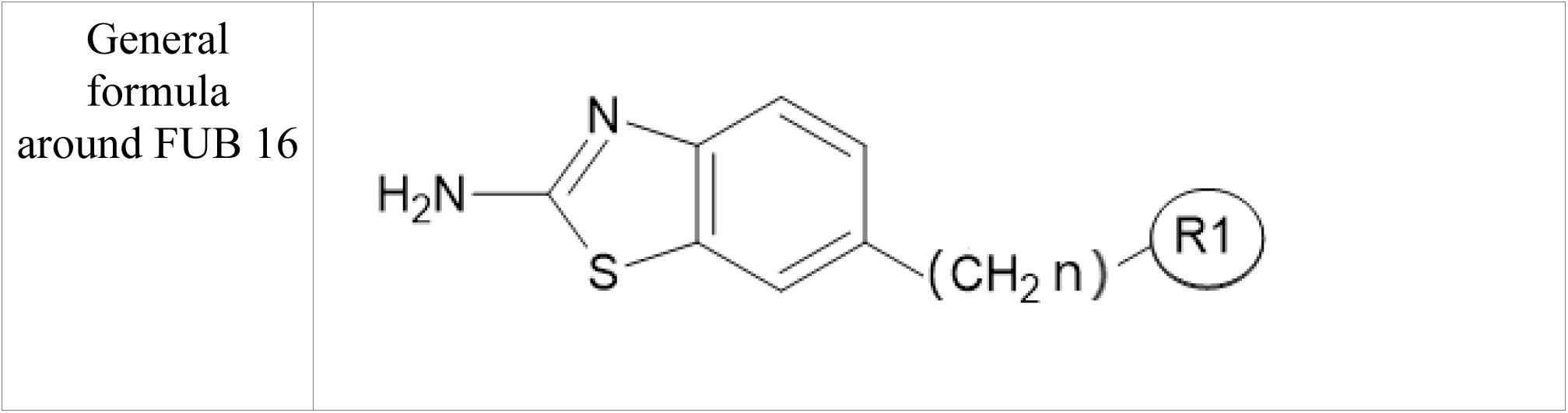

## THE ANTAGONIST

The tables give the IC 50 of the drugs in µM.

The first series describes antagonists, in which the molecules are derived from the general formula the first compound FUB 88 (derivated of R-(1H-imidazol-5-yl)ethan-1-amine) to the first interesting compound of this series FUB 114 (Table 10 to Table 12).

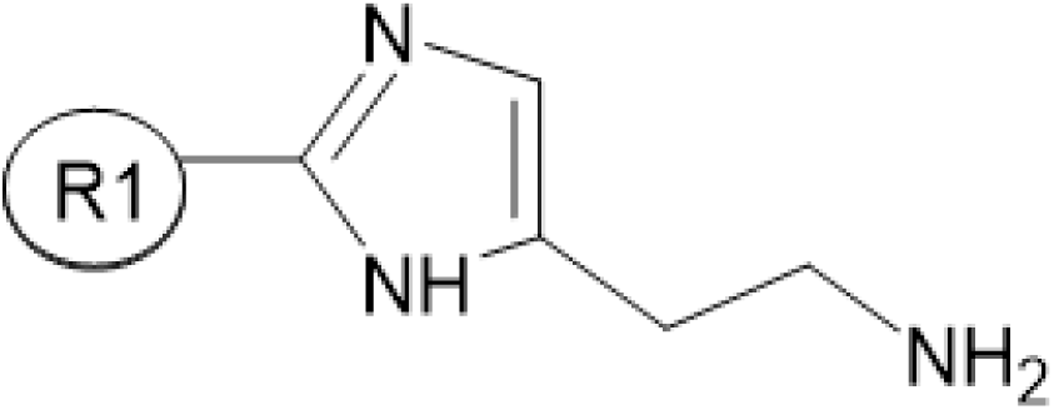

## DISCUSSION

Psychotic troubles are chronic and debilitating diseases with significant morbidity and mortality that often requires antipsychotic pharmacotherapy for life. Current therapy consists of neuroleptics with major anti-dopaminergic and anti-serotoninergic activity but may also act on other receptors such as histaminic and noradrenaline receptors (11).

It is well known from the prior art that typical neuroleptic agents induce extrapyramidal symptoms, which include rigidity, tremor, bradykinesia (slow movement) and bradyphrenia (slow though), as well as tardive dyskinesia, acute dystonic reactions and akathisia. Furthermore, atypical neuroleptic agents induce both extrapyramidal symptoms and other side effects such as increase of body weight, mood disturbance, sexual dysfunction, sedation, orthostatic hypotension, hypersalivation, lowered seizure threshold and, in particular, agranulocytosis (12)

Recent discoveries, brought to light the link between schizophrenia and bipolar disorders with disturbance in GABA and glutamate transmission in the brain. For example, schizophrenia would be associated with ionotropic N-methyl-D-aspartate (NMDA) receptor dysfunction (13). Indeed, according experimental researches, it has been found that NMDA receptor blockers such as phencyclidine (PCP) and MK-801 induce psychoses similar to that associated with schizophrenia (14,15). Since hypo function of NMDA system is considered to have an important role in schizophrenia and schizophreniform psychosis, especially negative symptoms, the fact that cognitive dysfunction caused by ketamine are similar to schizophrenia (16) reinforces this observation.

NMDA receptor modulators, such as antagonists, agonists and partial agonists have thus been the subject of several successive researches both for the treatment of psychotic diseases and for the treatment of central nervous diseases (17). For example, the NMDA receptor modulator memantine was developed for the treatment of Alzheimer’s disease (18). The partial agonist agent of D-cycloserine was revealed as having some antidepressant and anxiolytic activity (19). Furthermore, agents targeting the NMDA receptor appeared to be involved in different stages of development for the treatment of anxiety, depression, cognitive and motor disorders (20,21)

We have recently identified an histamine site in NMDA receptors. Histamine has numerous functions in the brain and in particular modulates responses of the NMDA receptors of hippocampal neurons (3). William K. (4) demonstrated that histamine could directly act at a novel recognition site on some subtypes of NMDA receptor to increase their activity. However, the histamine has shown a preferential effect on responses mediated by NR1/NR2B receptors (22).

The aim of this works was to provide compounds that can interfere with the NMDA receptor and in particular with the histamine site of the NMDA receptor. Recent studies also put light on the fact that distinct subtypes of the NMDA receptor are differently involved in central nervous system diseases. In particular, histamine site of the NMDA receptor may have a key role in several disorders. For example, NMDA receptor histamine site has been discovered and evidenced as being involved in Ischemia (23). We believe that enhancing NMDA receptor function will restore sensorimotor gating deficits observed in schizophrenia. Therefore, agonists of the NDMA receptor will be useful as anti-psychotic agents for the treatment of symptoms of this disease.

## CONCLUSION

In conclusion, these data confirm the existence of a histamine site, distinct from other allosteric sites, of the NMDAR. Since histamine also activates the human NMDAR (9), agonists of the NMDA_(HA)_R may be helpful in therapeutics. We suggest that agonists at the NMDA_(HA)_R constitute an innovative class of antipsychotics for the treatment of schizophrenia and other neurological or psychiatric disorders.

**This study was supported by INSERM, the French Ministère de la Recherche and the Fondation pour la Recherche Médicale**.

## AGONIST

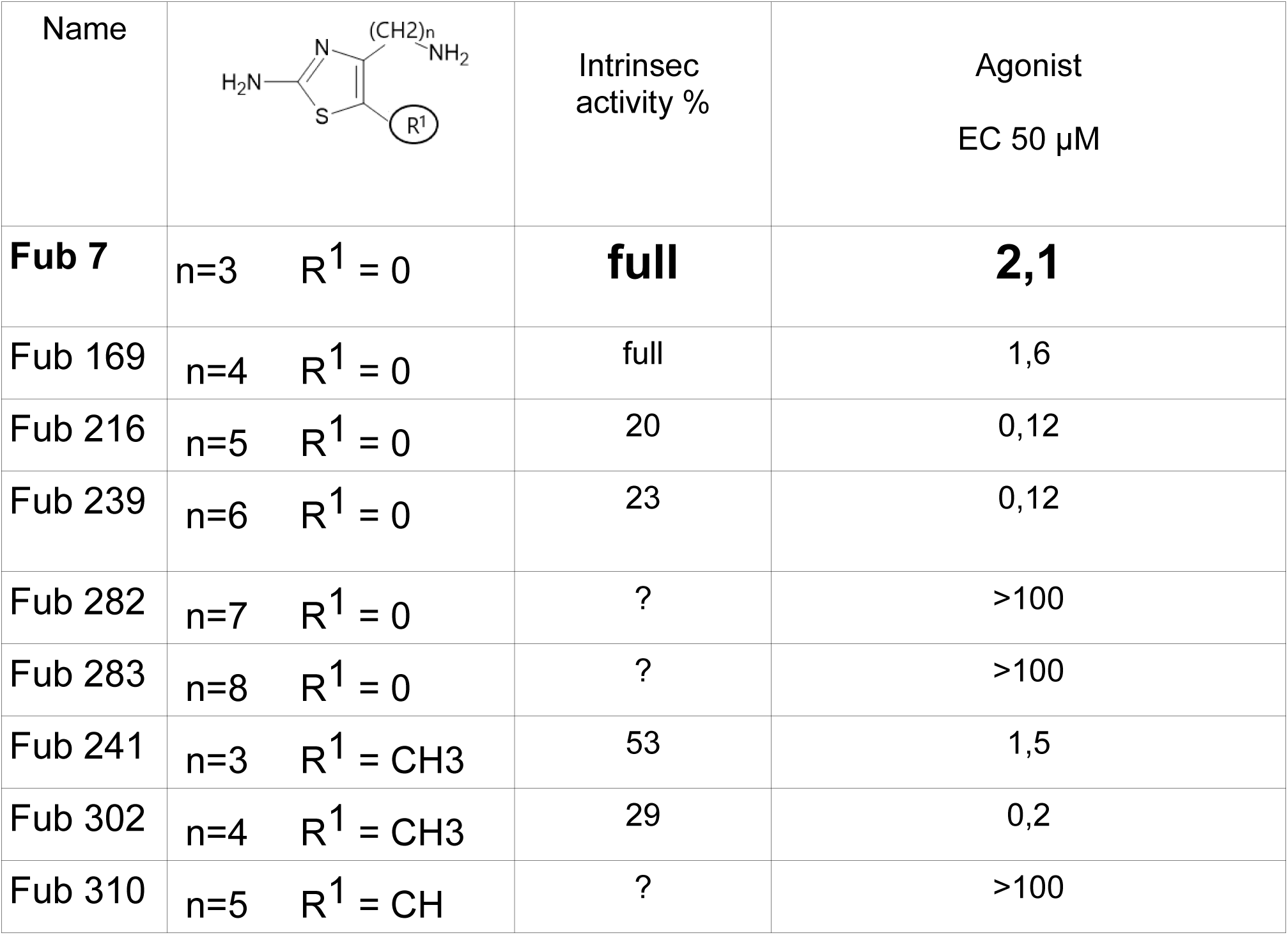
TABLE 1.

**TABLE 1**
| Name | | Intrinsic activity % | Agonist<br>EC 50 μM |
| --- | --- | --- | --- |
| **Fub 7** | n=3 R<sup>1</sup> = 0 | **full** | **2,1** |
| Fub 169 | n=4 R<sup>1</sup> = 0 | full | 1,6 |
| Fub 216 | n=5 R<sup>1</sup> = 0 | 20 | 0,12 |
| Fub 239 | n=6 R<sup>1</sup> = 0 | 23 | 0,12 |
| Fub 282 | n=7 R<sup>1</sup> = 0 | ? | >100 |
| Fub 283 | n=8 R<sup>1</sup> = 0 | ? | >100 |
| Fub 241 | n=3 R<sup>1</sup> = CH<sub>3</sub> | 53 | 1,5 |
| Fub 302 | n=4 R<sup>1</sup> = CH<sub>3</sub> | 29 | 0,2 |
| Fub 310 | n=5 R<sup>1</sup> = CH | ? | >100 |

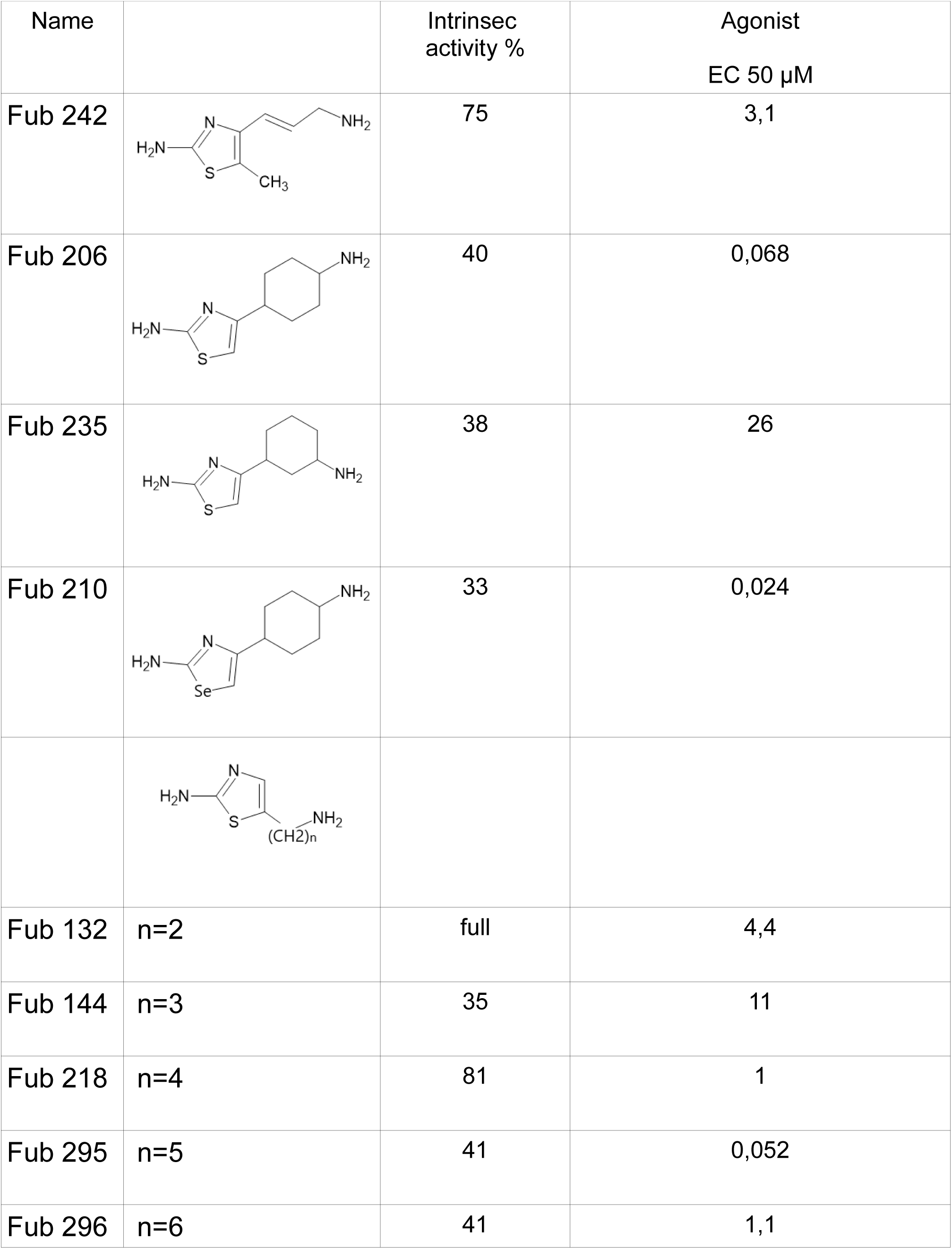
TABLE 2.

**TABLE 2**
| Name | | Intrinsec activity % | Agonist<br>EC 50 μM |
| --- | --- | --- | --- |
| Fub 242 |  | 75 | 3,1 |
| Fub 206 |  | 40 | 0,068 |
| Fub 235 |  | 38 | 26 |
| Fub 210 |  | 33 | 0,024 |
| Fub 132 | n=2 | full | 4,4 |
| Fub 144 | n=3 | 35 | 11 |
| Fub 218 | n=4 | 81 | 1 |
| Fub 295 | n=5 | 41 | 0,052 |
| Fub 296 | n=6 | 41 | 1,1 |

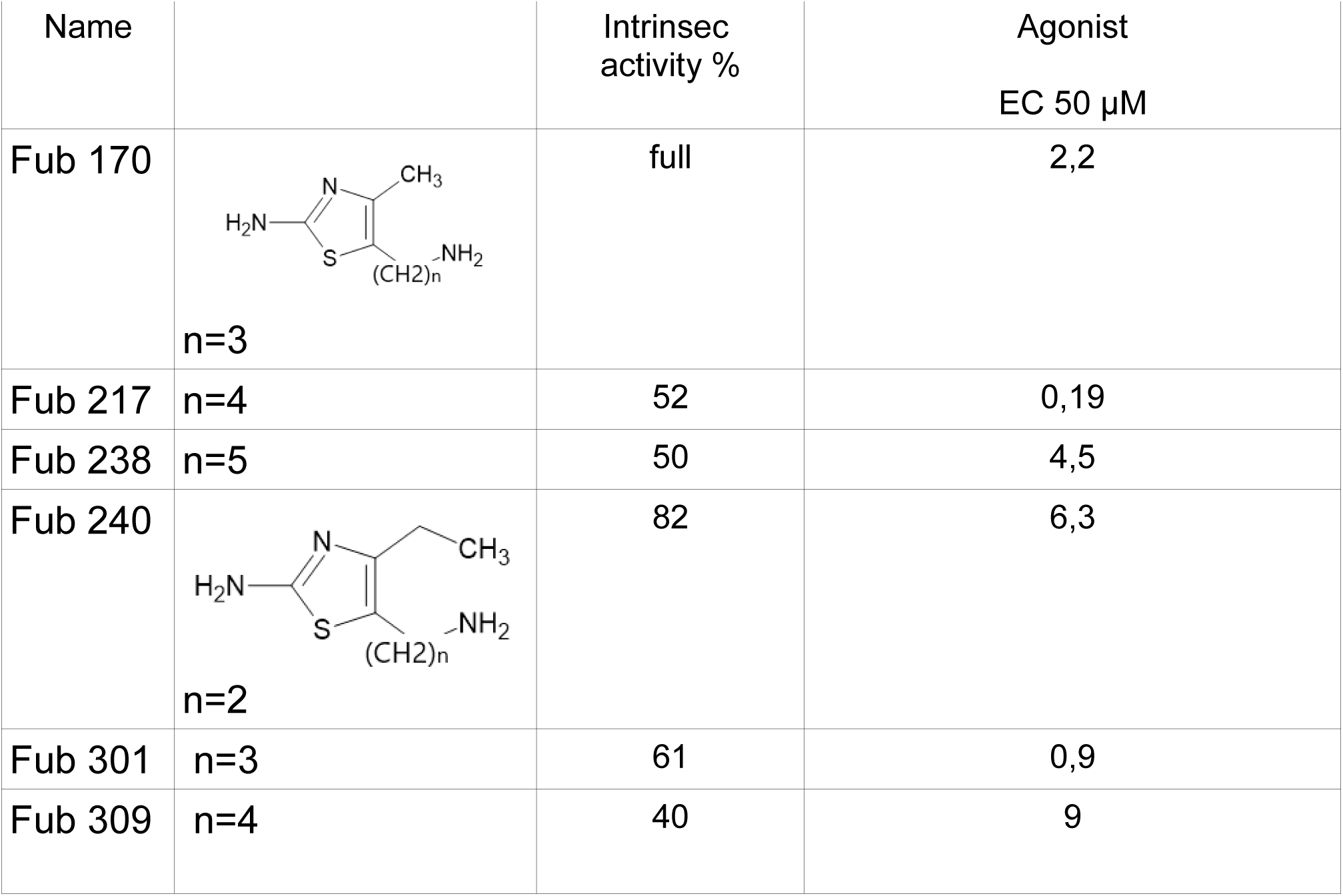
TABLE 3.

**TABLE 3**
| Name | | Intrinsec<br>activity % | Agonist<br>EC 50 μM |
| --- | --- | --- | --- |
| Fub 170 | n=3 | full | 2,2 |
| Fub 217 | n=4 | 52 | 0,19 |
| Fub 238 | n=5 | 50 | 4,5 |
| Fub 240 | n=2 | 82 | 6,3 |
| Fub 301 | n=3 | 61 | 0,9 |
| Fub 309 | n=4 | 40 | 9 |

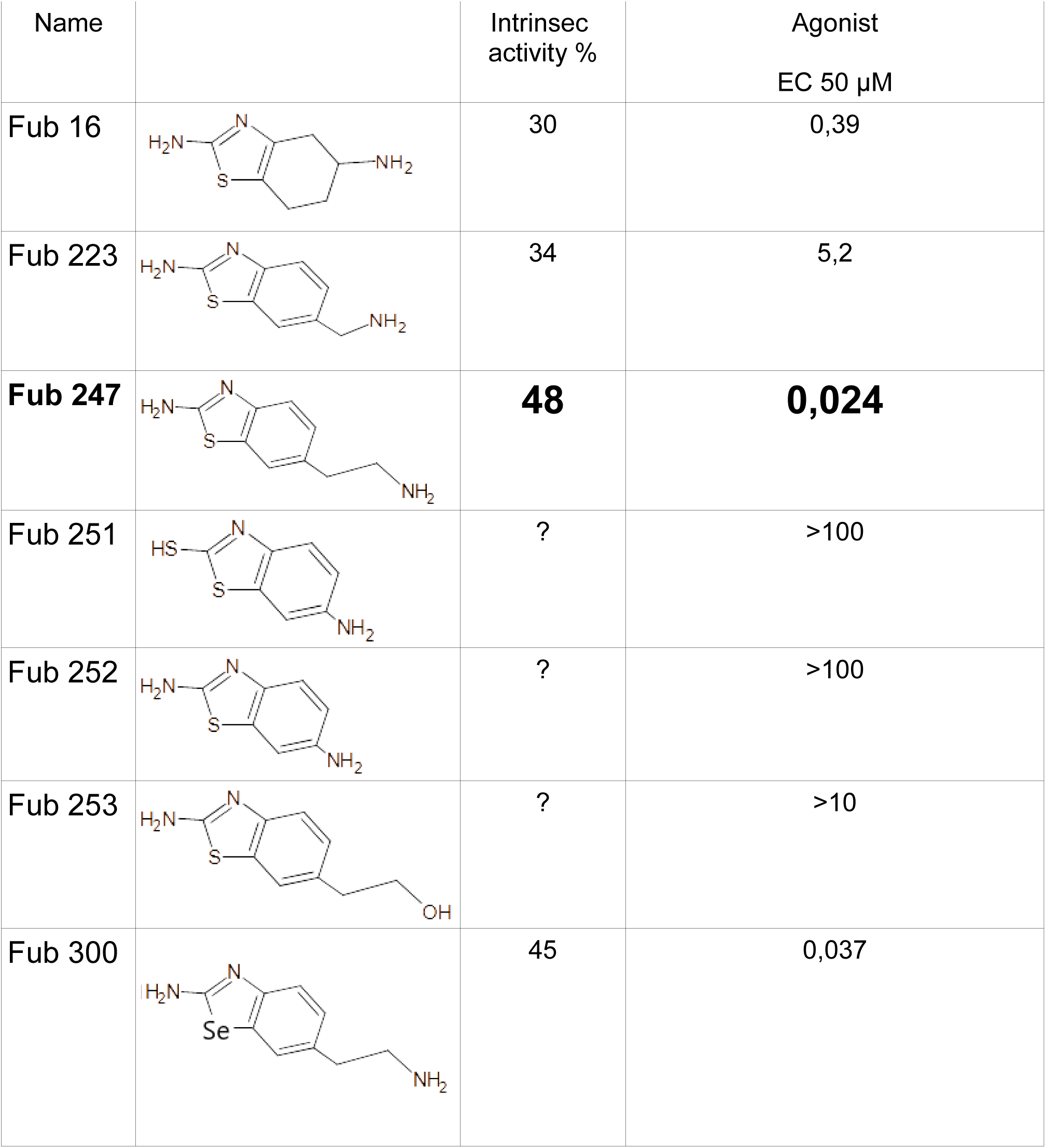
TABLE 4.

**TABLE 4**
| Name | | Intrinsec activity % | Agonist<br>EC 50 μM |
| --- | --- | --- | --- |
| Fub 16 |  | 30 | 0,39 |
| Fub 223 |  | 34 | 5,2 |
| **Fub 247** |  | **48** | **0,024** |
| Fub 251 |  | ? | >100 |
| Fub 252 |  | ? | >100 |
| Fub 253 |  | ? | >10 |
| Fub 300 |  | 45 | 0,037 |

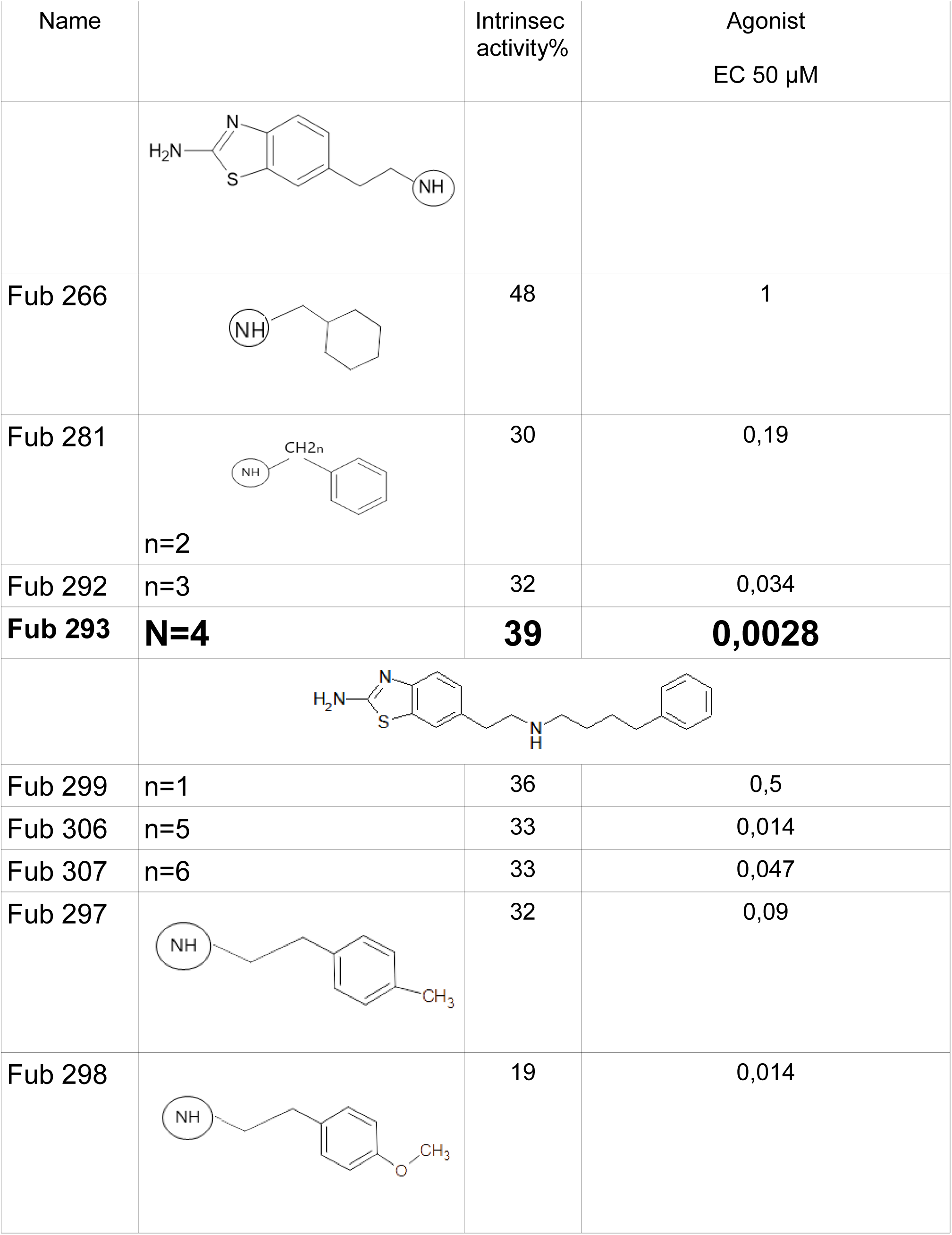
TABLE 5.

**TABLE 5**
| Name | | Intrinsec activity% | Agonist<br>EC 50 μM |
| --- | --- | --- | --- |
| Fub 266 |  | 48 | 1 |
| Fub 281 | n=2 | 30 | 0,19 |
| Fub 292 | n=3 | 32 | 0,034 |
| **Fub 293** | **N=4** | **39** | **0,0028** |
| Fub 299 | n=1 | 36 | 0,5 |
| Fub 306 | n=5 | 33 | 0,014 |
| Fub 307 | n=6 | 33 | 0,047 |
| Fub 297 |  | 32 | 0,09 |
| Fub 298 |  | 19 | 0,014 |

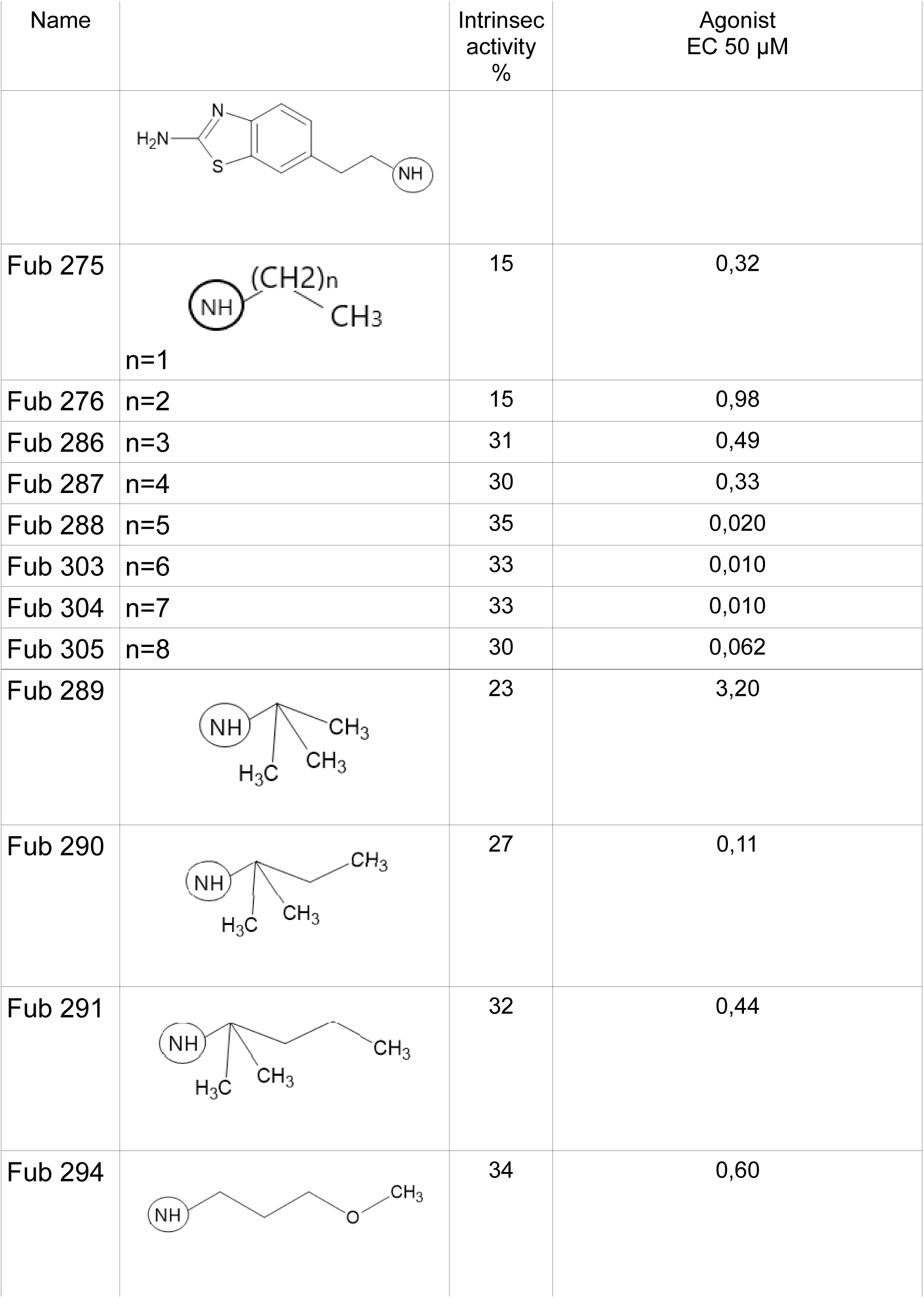
TABLE 6.

**TABLE 6**
| Name | | Intrinsec activity % | Agonist EC 50 μM |
| --- | --- | --- | --- |
| Fub 275 | n=1 | 15 | 0,32 |
| Fub 276 | n=2 | 15 | 0,98 |
| Fub 286 | n=3 | 31 | 0,49 |
| Fub 287 | n=4 | 30 | 0,33 |
| Fub 288 | n=5 | 35 | 0,020 |
| Fub 303 | n=6 | 33 | 0,010 |
| Fub 304 | n=7 | 33 | 0,010 |
| Fub 305 | n=8 | 30 | 0,062 |
| Fub 289 |  | 23 | 3,20 |
| Fub 290 |  | 27 | 0,11 |
| Fub 291 |  | 32 | 0,44 |
| Fub 294 |  | 34 | 0,60 |

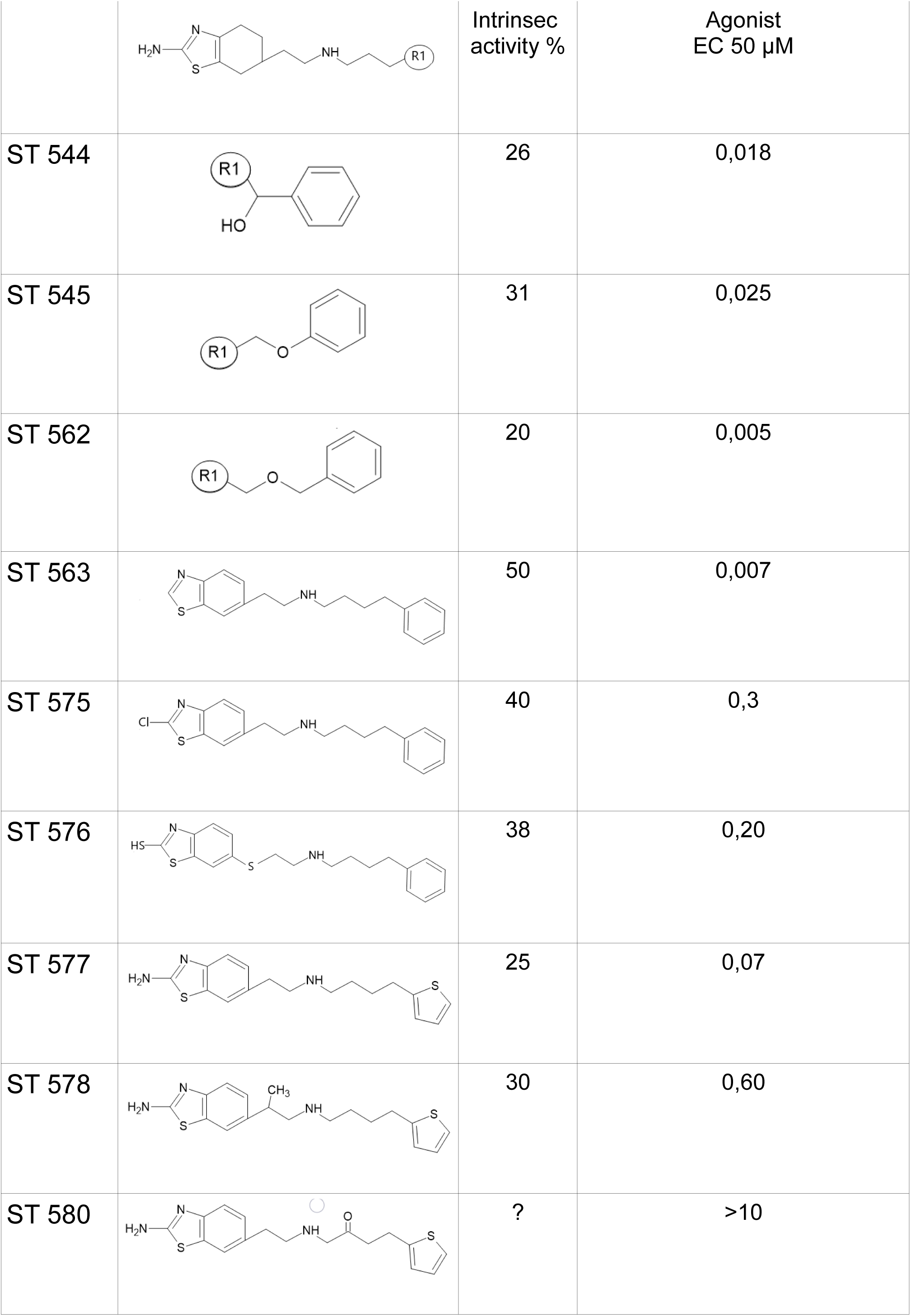
TABLE 7.

**TABLE 7**
|  |  |  |  |
| --- | --- | --- | --- |
| | | Intrinsic activity % | Agonist EC 50 μM |
| ST 544 |  | 26 | 0,018 |
| ST 545 |  | 31 | 0,025 |
| ST 562 |  | 20 | 0,005 |
| ST 563 |  | 50 | 0,007 |
| ST 575 |  | 40 | 0,3 |
| ST 576 |  | 38 | 0,20 |
| ST 577 |  | 25 | 0,07 |
| ST 578 |  | 30 | 0,60 |
| ST 580 |  | ? | >10 |

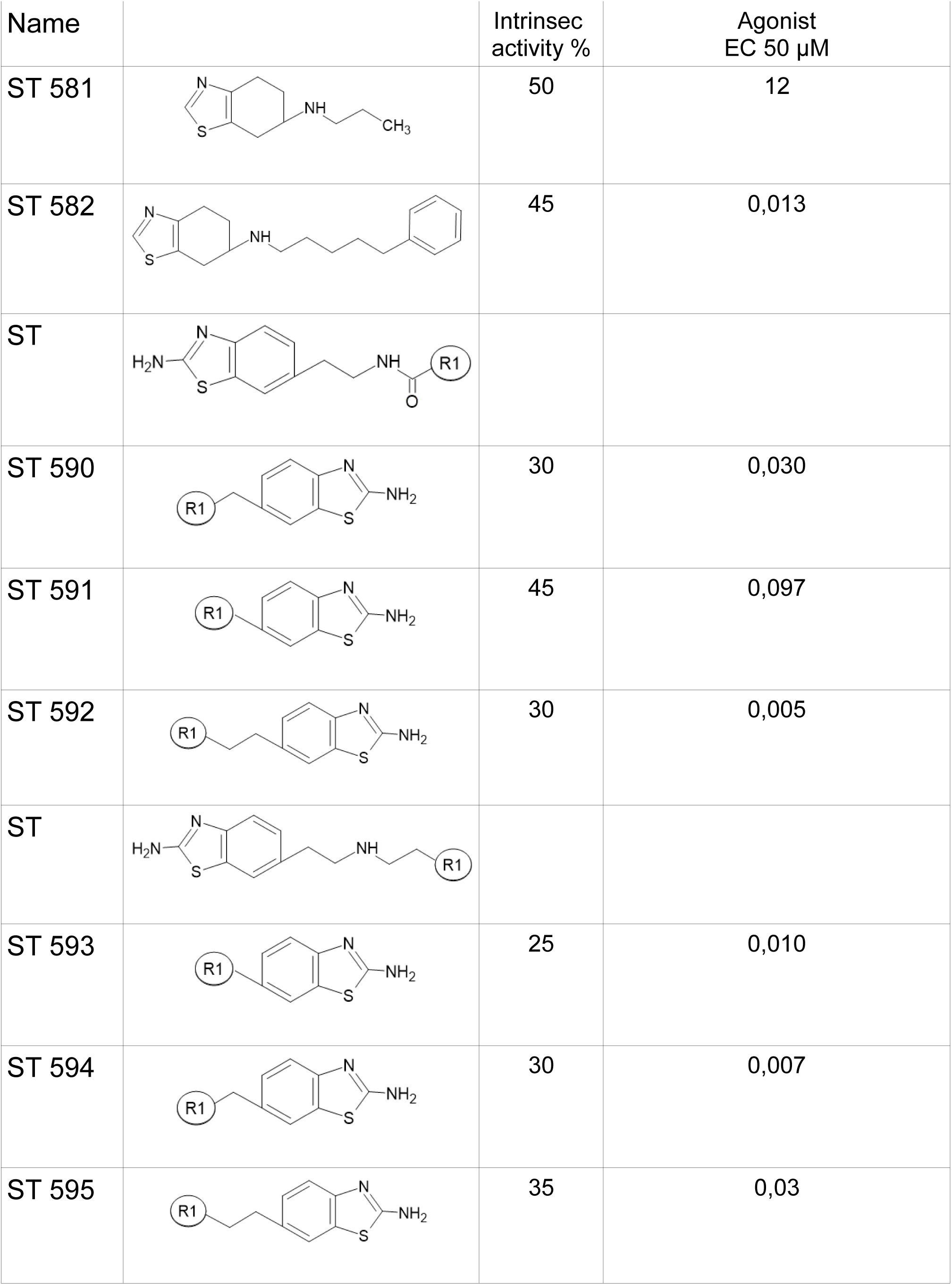
TABLE 8.

**TABLE 8**
| Name | | Intrinsec activity % | Agonist EC 50 μM |
| --- | --- | --- | --- |
| ST 581 |  | 50 | 12 |
| ST 582 |  | 45 | 0,013 |
| ST |  |  |  |
| ST 590 |  | 30 | 0,030 |
| ST 591 |  | 45 | 0,097 |
| ST 592 |  | 30 | 0,005 |
| ST |  |  |  |
| ST 593 |  | 25 | 0,010 |
| ST 594 |  | 30 | 0,007 |
| ST 595 |  | 35 | 0,03 |

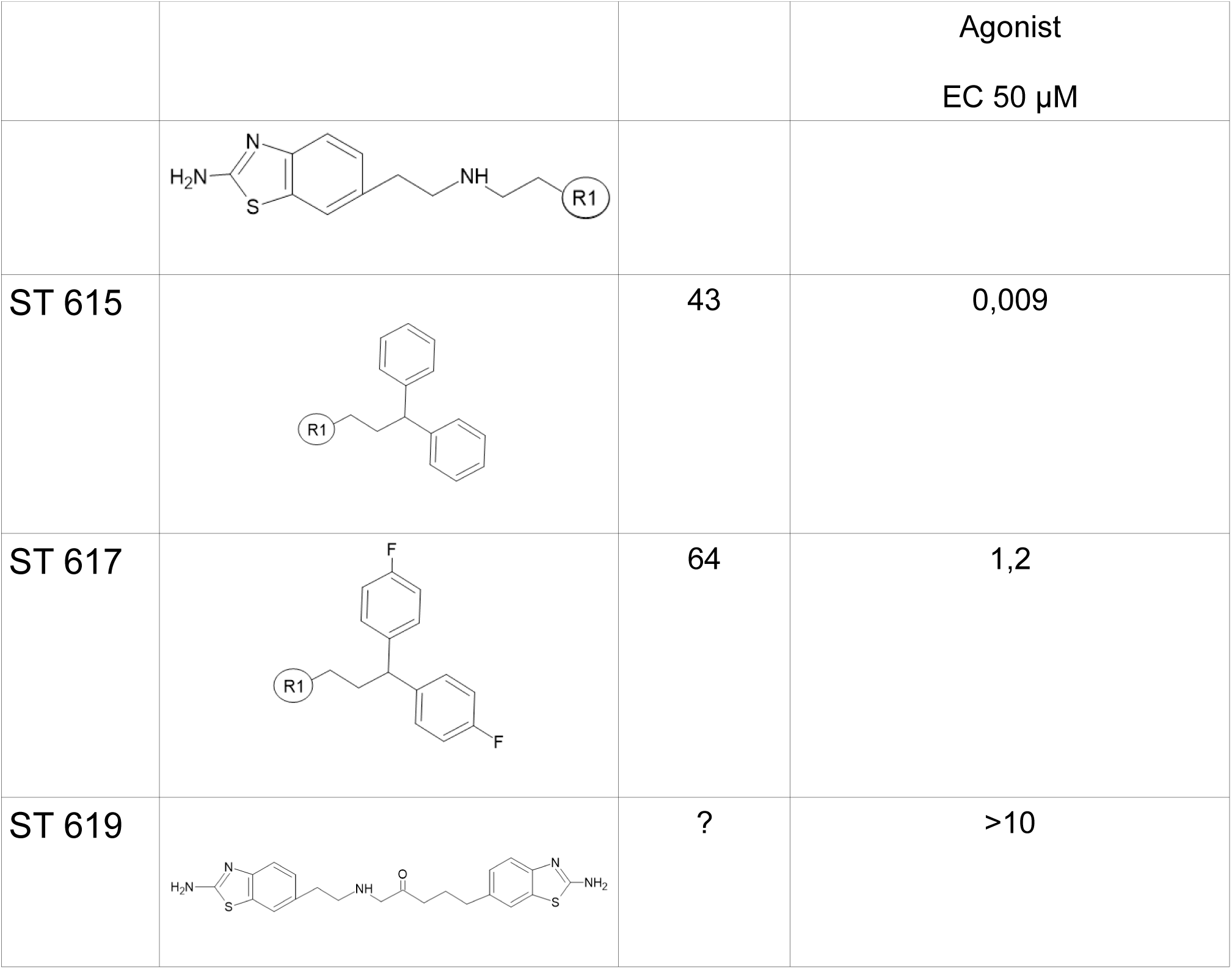
TABLE 9.

**TABLE 9**
| | | | Agonist<br>EC 50 μM |
| --- | --- | --- | --- |
| ST 615 |  | 43 | 0,009 |
| ST 617 |  | 64 | 1,2 |
| ST 619 |  | ? | >10 |

## ANTAGONIST

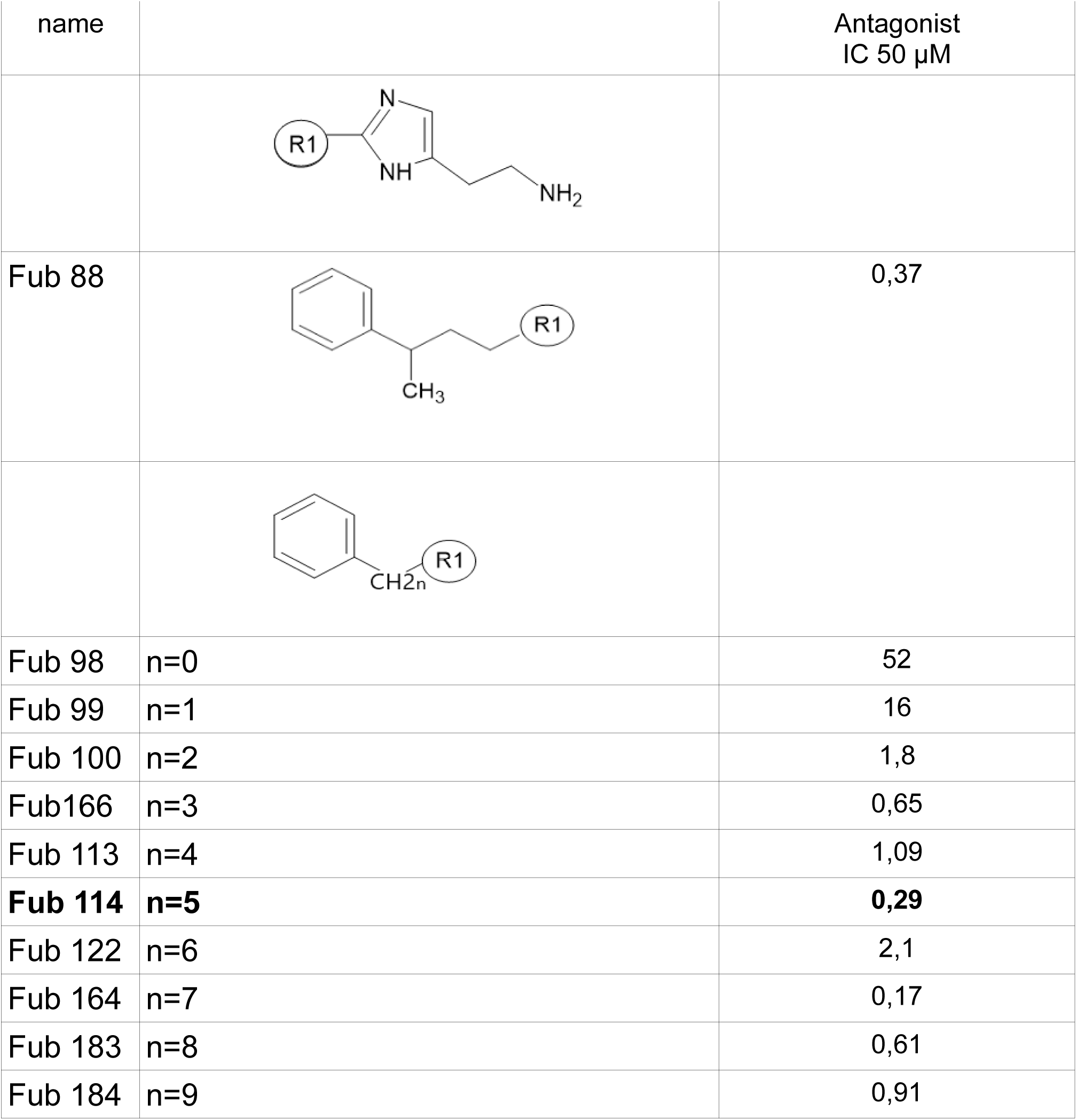
TABLE 10.

TABLE 10
| name | | Antagonist<br>IC 50 μM |
| --- | --- | --- |
| Fub 88 |  | 0,37 |
| Fub 98 | n=0 | 52 |
| Fub 99 | n=1 | 16 |
| Fub 100 | n=2 | 1,8 |
| Fub166 | n=3 | 0,65 |
| Fub 113 | n=4 | 1,09 |
| **Fub 114** | **n=5** | **0,29** |
| Fub 122 | n=6 | 2,1 |
| Fub 164 | n=7 | 0,17 |
| Fub 183 | n=8 | 0,61 |
| Fub 184 | n=9 | 0,91 |

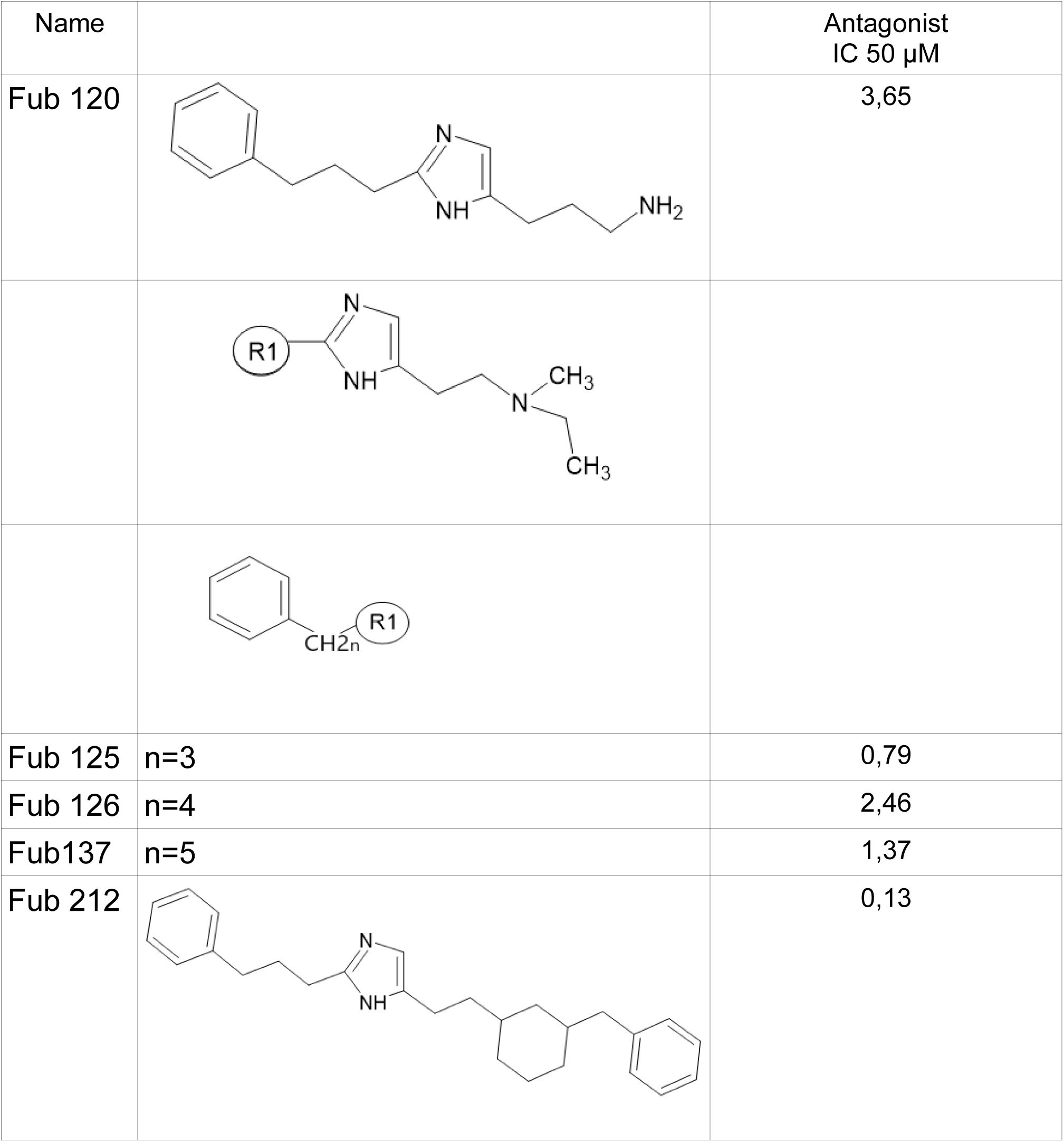
TABLE 11.

**TABLE 11**
| Name | | Antagonist<br>IC 50 μM |
| --- | --- | --- |
| Fub 120 |  | 3,65 |
| Fub 125 | n=3 | 0,79 |
| Fub 126 | n=4 | 2,46 |
| Fub137 | n=5 | 1,37 |
| Fub 212 |  | 0,13 |

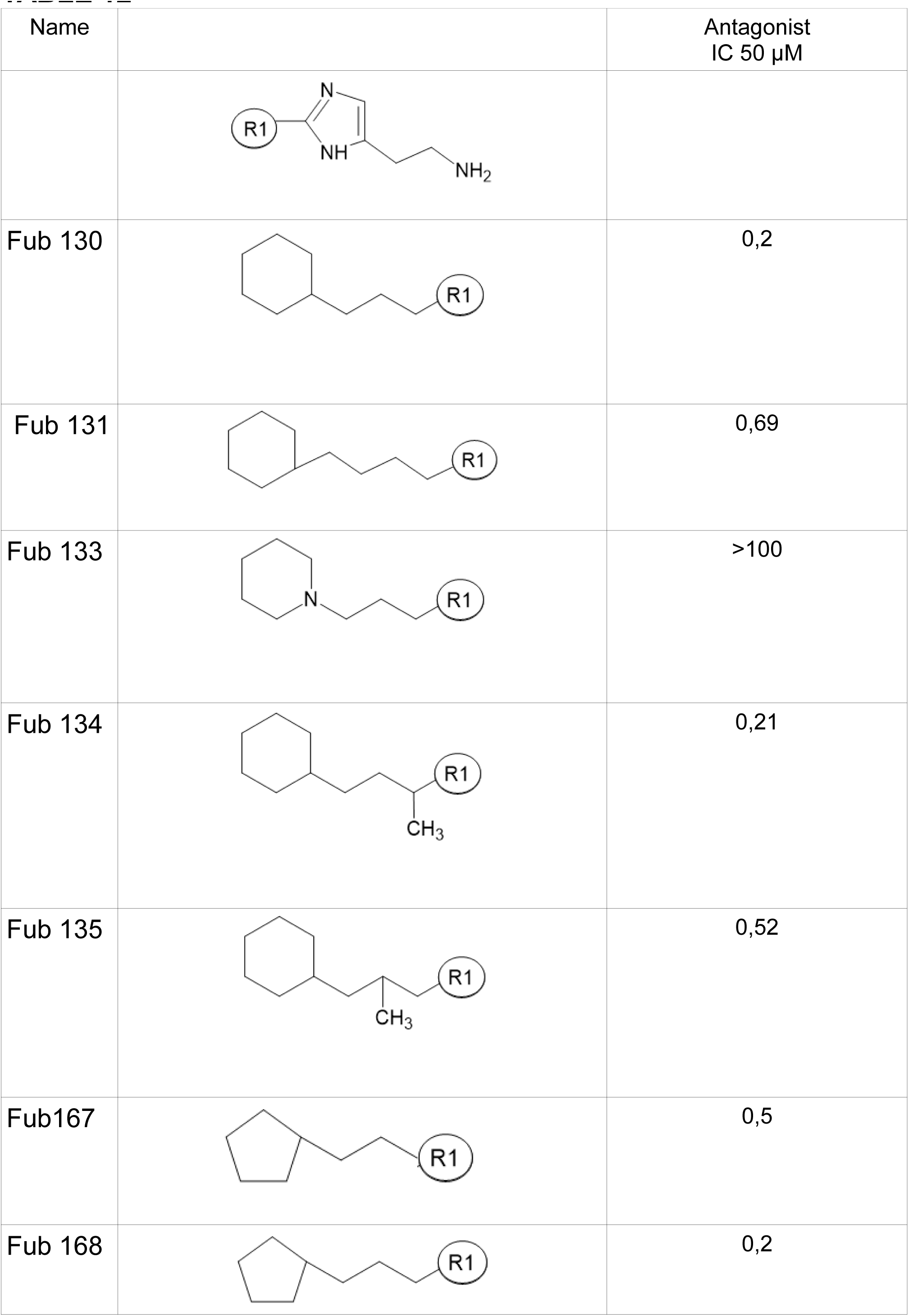
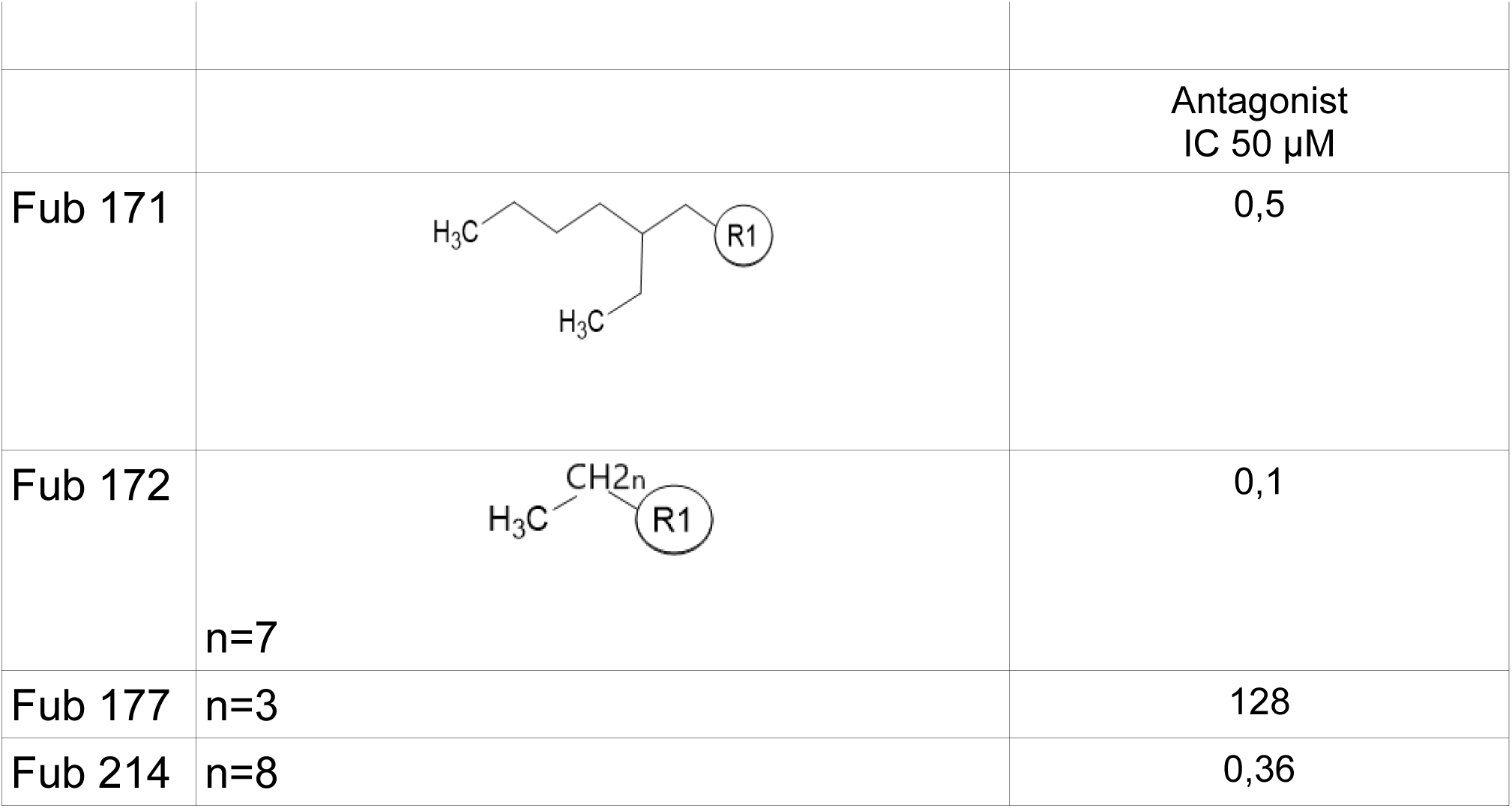
TABLE 12.

**TABLE 12**
| Name | | Antagonist<br>IC 50 μM |
| --- | --- | --- |
| Fub 130 |  | 0,2 |
| Fub 131 |  | 0,69 |
| Fub 133 |  | >100 |
| Fub 134 |  | 0,21 |
| Fub 135 |  | 0,52 |
| Fub167 |  | 0,5 |
| Fub 168 |  | 0,2 |
| | | Antagonist<br>IC 50 μM |
| Fub 171 |  | 0,5 |
| Fub 172 | n=7 | 0,1 |
| Fub 177 | n=3 | 128 |
| Fub 214 | n=8 | 0,36 |

The second series describes the antagonist, in which the molecules are derived from the general formula around the lead compound FUB 16 and lead to ST-579 the most potent drug realized (Table 13 and Table 14).

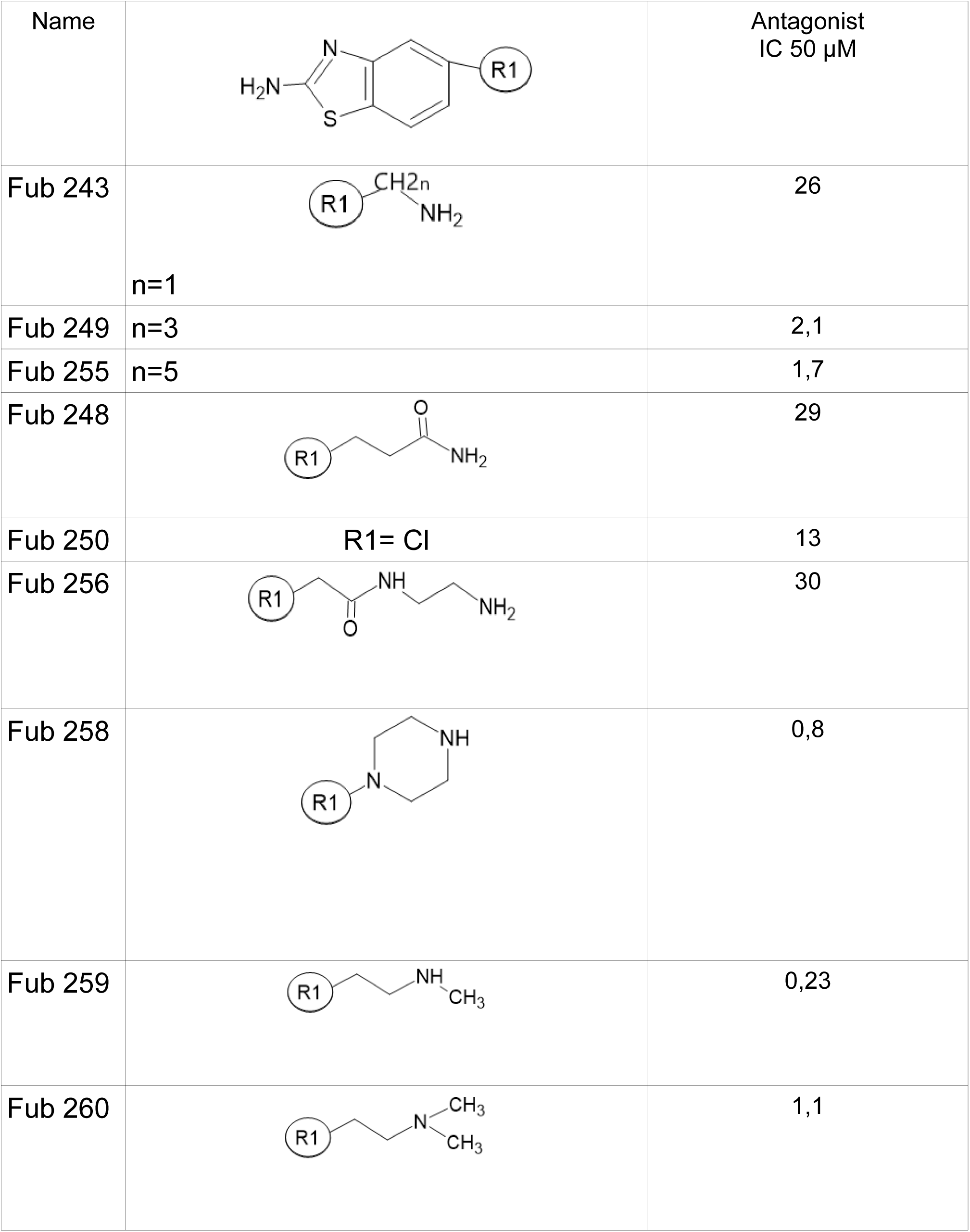
TABLE 13.

**TABLE 13**
| Name | | Antagonist<br>IC 50 μM |
| --- | --- | --- |
| Fub 243 | n=1 | 26 |
| Fub 249 | n=3 | 2,1 |
| Fub 255 | n=5 | 1,7 |
| Fub 248 |  | 29 |
| Fub 250 | R1= Cl | 13 |
| Fub 256 |  | 30 |
| Fub 258 |  | 0,8 |
| Fub 259 |  | 0,23 |
| Fub 260 |  | 1,1 |

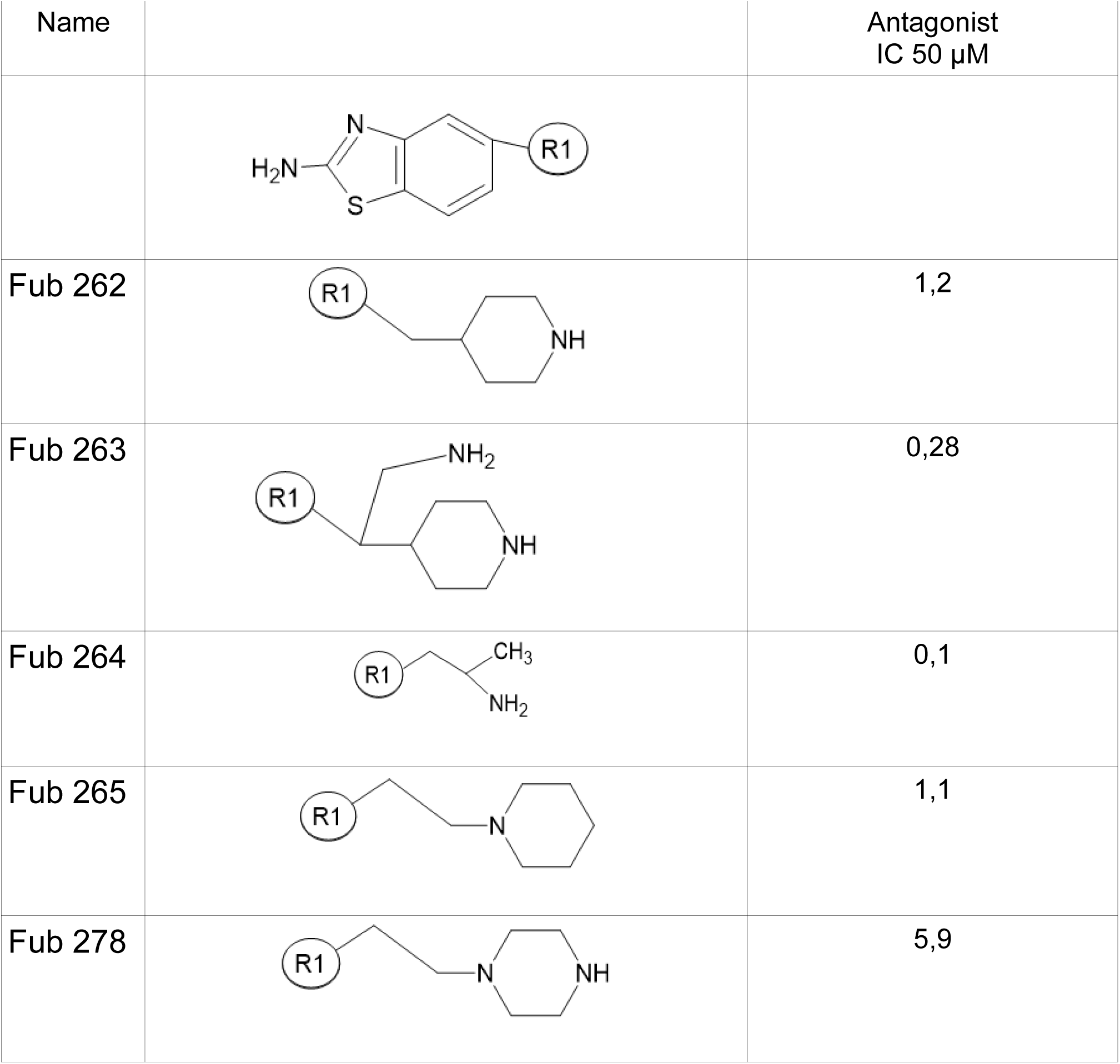
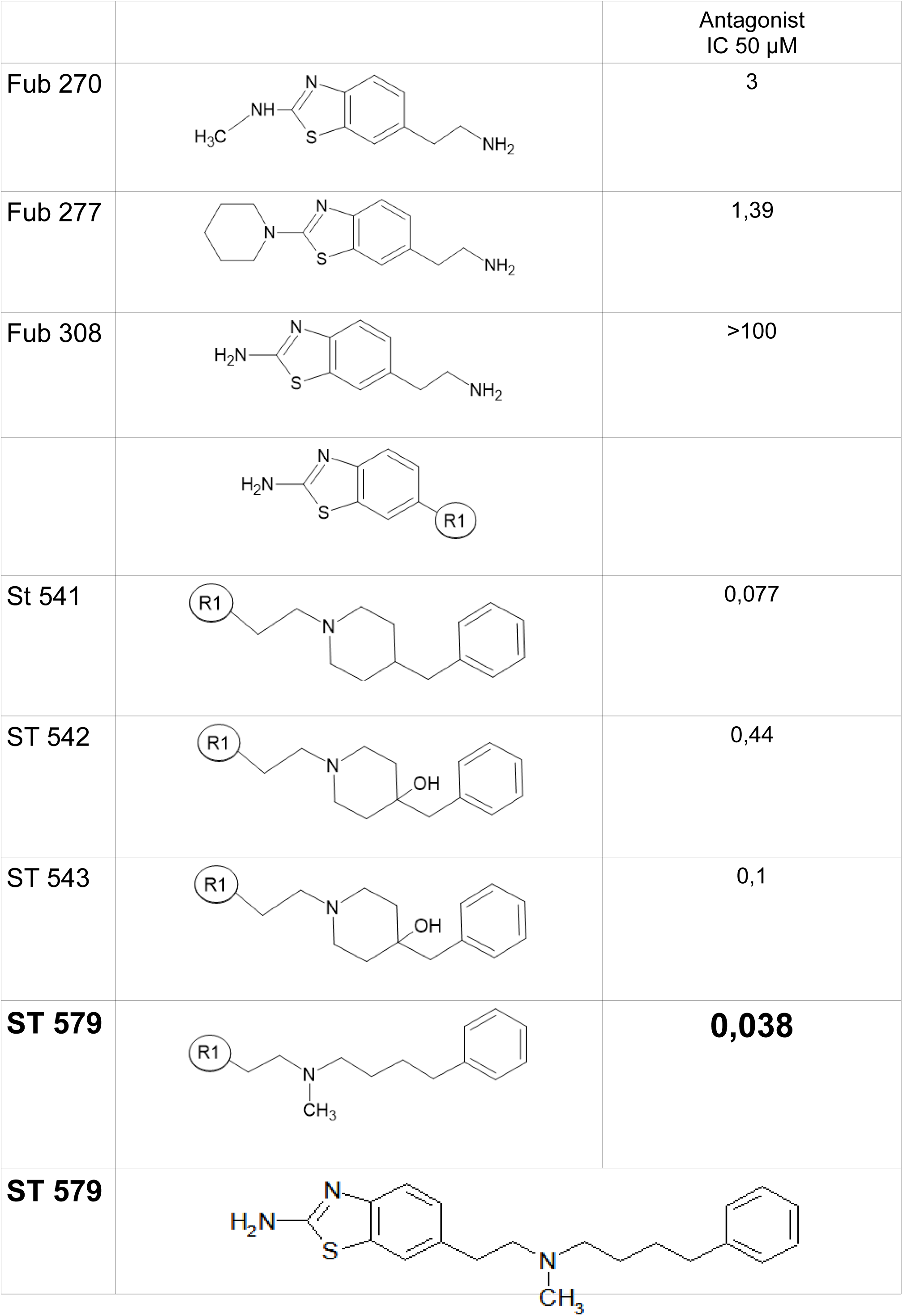
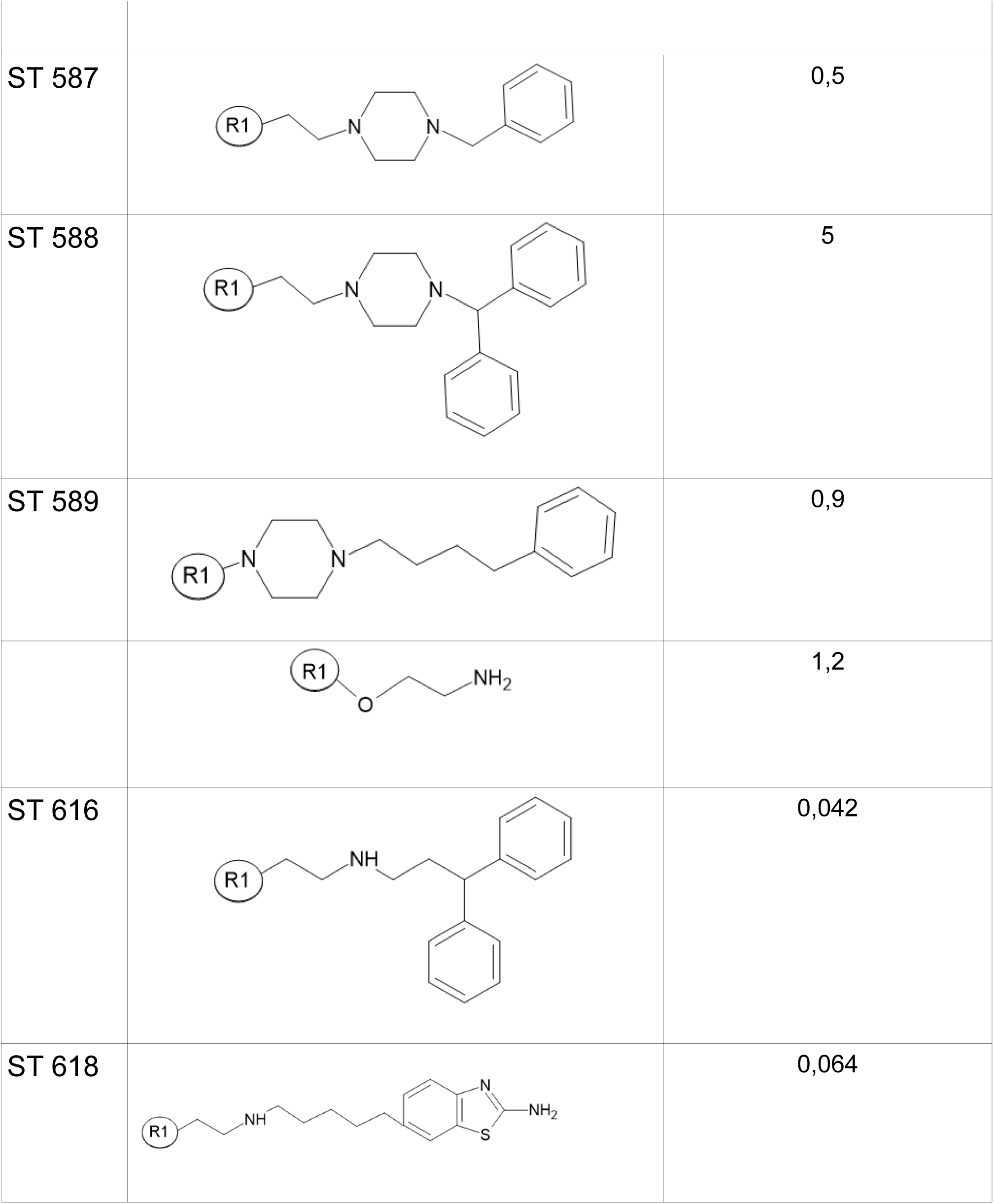
TABLE 14.

**TABLE 14**
| Name | | Antagonist<br>IC 50 μM |
| --- | --- | --- |
| Fub 262 |  | 1,2 |
| Fub 263 |  | 0,28 |
| Fub 264 |  | 0,1 |
| Fub 265 |  | 1,1 |
| Fub 278 |  | 5,9 |

| | | Antagonist<br>IC 50 μM |
| --- | --- | --- |
| Fub 270 |  | 3 |
| Fub 277 |  | 1,39 |
| Fub 308 |  | >100 |
| St 541 |  | 0,077 |
| ST 542 |  | 0,44 |
| ST 543 |  | 0,1 |
| **ST 579** |  | **0,038** |
| **ST 579** |  |  |
| ST 587 |  | 0,5 |
| ST 588 |  | 5 |
| ST 589 |  | 0,9 |
|  |  | 1,2 |
| ST 616 |  | 0,042 |
| ST 618 |  | 0,064 |

